# Salt leverages polyethylene terephthalate hydrolase (PETase) enzymatic activity via the predicted conformational switch

**DOI:** 10.1101/2021.09.23.461413

**Authors:** Sergey A. Shiryaev, Piotr Cieplak, Andrei V. Chernov

## Abstract

Plastic pollution spawned a global challenge caused by the environmental accumulation of polyethylene terephthalate (PET) plastics. Ongoing remediation efforts using microbial and engineered PET hydrolyzing enzymes (PETases) are hindered by slow depolymerization activities. Here, we report the optimized reaction conditions that leveraged the PETase hydrolase activity 2 to 3.8-fold in the presence of high NaCl concentrations. Molecular dynamics simulations (MDS) were applied to model salt-dependent conformational changes of the PETase enzyme bound to a 3-unit PET polymer. MDS demonstrated that PETase interaction with flanking polymer units exhibited a striking structural disparity at low and high salt concentrations. At low salt concentrations, flanking polymer units displayed significant bending. In contrast, flanking units extended at high salt concentrations, thus residues D206, H237, and S160 of the catalytic triad positioned in close vicinity of the scissile ester bond of the polymer substrate. The resulting high salt-specific enzyme/substrate geometry can potentially facilitate hydrolysis. We theorized that a salt-dependent conformational switch could attenuate the enzyme to a broad range of natural and artificial polymers consumed as carbon sources. Altogether, new knowledge may advance the engineering of PET hydrolase enzymes and benefit bioconversion programs.

## Introduction

Polyethylene terephthalate (PET) is a versatile plastic mass-produced for a variety of industrial and consumer products. The annual worldwide manufacturing of PET has reached 50 million tons and is expected to double every five years. Over 30% of post-consumer plastic waste is amounted by PET. Although PET waste recycling has improved, accelerating industrial production and high resistance to environmental degradation impact recycling efforts. Especially concerning are PET microparticles that accumulate in ocean water and severely impact marine life (1, 2).

Bioconversion and bioremediation are two ways to reduce the adverse effects of plastic pollution. In response to the environmental impact of accumulating plastic waste, specialized enzymatic and catabolic pathways that utilize plastics as a carbon and energy source have naturally evolved. Bacteria, yeast, and fungi inhabiting environments with plastic debris can potentially use PET as a primary carbon source. *Ideonella sakaiensis*, a bacteria isolated at a recycling facility, can decompose PET, albeit relatively slowly (3). *I. sakaiensis* secretes two PET degrading enzymes: PETase and MHETase. PETase hydrolyses PET polymers to mono-2-hydroxyethyl (MHET) and bis-2-hydroxyethyl terephthalic acids (BHET). MHETase further converts MHET into ethylene glycol that breaks down in soil (4). An ongoing effort by the international community aimed to re-engineer enzymes with increased activity and stability for *in situ* remediations (5–16). However, engineered enzymes that exhibit high enzymatic activity under varying and challenging environmental conditions, including high or low temperatures, extreme acidity, or high salinity, will be required.

The optimization of PETase enzymatic characteristics that drastically increased the hydrolytic activity is reported here. We combined biochemical assays and molecular dynamic simulations (MDS) to propose a model of enzyme-substrate interaction under optimized conditions.

## Materials and methods

### Reagents

p-Nitrophenyl acetate (p-NPA) was from TCI America (Portland, OR, USA). Horseradish peroxidase (HRP)-conjugated donkey anti-mouse IgGs was from Jackson ImmunoResearch Laboratories (West Grove, PA, USA). All other reagents were from MilliporeSigma (Burlington, MA, USA)

### Enzyme purification

cDNA sequence of PETase was derived from *Ideonella sakaiensis* isolate amino acid sequence (3) with modifications described (12). Flag-tag and 9xHistidine tag were added at the C-terminus of the PETase. Synthetic constructs were manufactured by Genscript (Piscataway, NJ, USA) and inserted into the pET-21b(+) vector (Novogene, Boston, MA, USA) at *Nde*I and *Xho*I sites to generate pET-21b-PETase-SW expression plasmid.

Protein expression was conducted in *E.coli* BL21 (DE3) Codon Plus cells (Agilent Technologies, San Diego, CA, USA). Induced cultures were grown at 18° C overnight. Cells were collected by centrifugation (5,000g for 20 min), and pellets were resuspended in 50 ml of purification buffer (Tris-HCl buffer, pH 8.0, 500 mM NaCl supplemented with protease inhibitors). Cell suspensions were incubated with 5 mg/ml lysozyme on ice for 30 min and disrupted by sonication. Proteins were purified using Protino NTA columns (Macherey-Nagel, Freemansburg, PA, USA) using linear imidazole gradients (30-250 mM). Protein identity was confirmed by immunoblotting. Protein concentrations were measured using Coomassie Protein Assays. Purified proteins were concentrated by dialysis in storage buffer (20 mM Tris-HCl buffer, pH 8.0, 500 mM NaCl supplemented with 50% glycerol). Aliquots were stored at −80° C.

### PETase activity assay

PETase activity was measured in 96 wells plates for 20 min by optical absorption at 400 nm using SpectraFluor Plus microplate reader (Tecan US, Morrisville, NC, USA). Reactions were conducted in 200 μl of the reaction buffer (20 mM Tris-HCl, 100 mM NaCl, 0.005% Brij 35) supplemented with 100 nM PETase. To initiate the chromogenic reaction, 100 μM p-NPA was added to each well. All enzymatic activities were measured at 24° C and pH 8.0 unless indicated overwise. Optimization parameters included pH (4.0-9.0), temperature (24-50° C), glycerol (0–25%), NaCl (0.1–5M), SDS (0–4%) and DMSO (0-100%) that were tested in respective increments as indicated on Fig. 1. Fold-change of the specific activity was calculated relative to the baseline. To control the non-enzymatic hydrolysis of p-NPA, parallel reactions for each optimization parameter were conducted without PETase.

**Figure 1.**
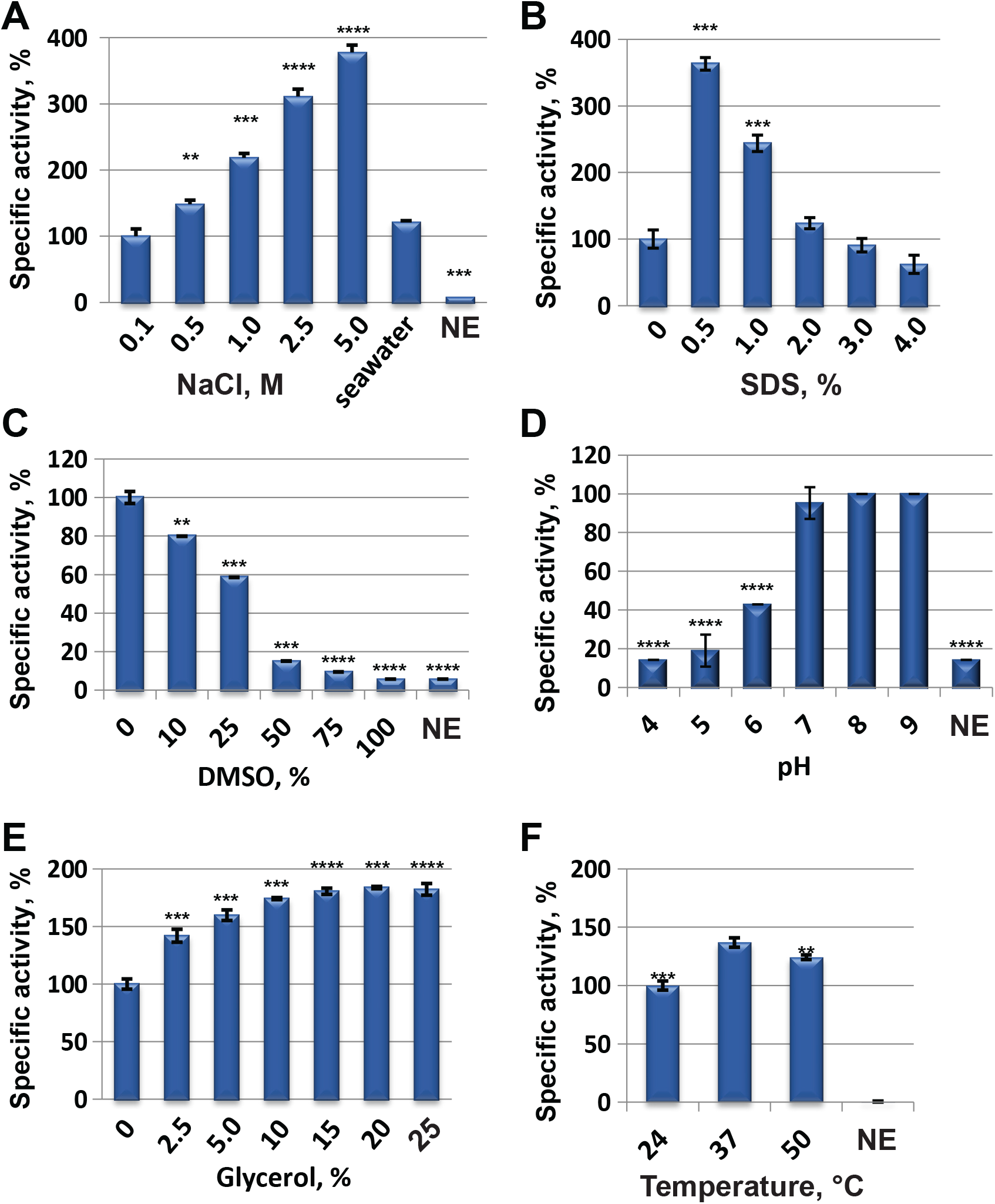
Optimization of PETase enzymatic activity. **(A)** PETase activity is stimulated by a gradual increase of NaCl (0.1-5M); **(B)** SDS; **(C)** DMSO; **(D)** pH; **(E)** glycerol; **(F)** temperature. NE, no enzyme. A two-tailed unpaired t-test with Welch’s parametric test was used. * - p<0.05, ** - p<0.005, *** - p<0.0005 and **** - p<0.0001. Baseline activity (100%) was measured in 20 mM Tris-HCl, pH 8.0, 100 mM NaCl, 0.005% Brij 35.

### Statistical and kinetics analysis

A two-tailed unpaired t-test with Welch’s correlations parametric test was used to compare enzyme activities between test groups. P-value <0.05 was considered significant. Enzymatic parameters were calculated and visualized using GraphPad Prism (GraphPad Software Inc, USA).

### MD simulations

3D structures of PETase (PDB:6ANE) (11) and 1-2(-hydroxyethyl) 4-methyl terephthalate (HEMT) (PDB:5XH3) (17) were from Protein Data Bank (https://www.rcsb.org). Molecular dynamics (MD) simulations with and without docked substrate were performed using the AMBER18 package (18, 19). The systems were placed in the truncated octahedron periodic box of the TIP3P water model (20). The distance between protein and the edge of the box to at least 1 nm. Electrostatic interactions were treated using Particle Mesh Ewald (PME) approach (21) with a 0.16 nm grid, fourth-order cubic interpolations, and tolerance of 10^−5^. Van der Waals interactions were calculated using a 10 Å cutoff. Simulations systems were minimized using the steepest descent method, then the systems were gradually heated to 300K and equilibrated. Production runs were performed using NPT ensembles (p=1 atm, T=300K, 0.5 μsec timescale). FF14SB (22) and GAFF force field parameters (23) were applied to the protein and substrate model, respectively. Atomic charges were determined using the RESP model (24). Simulations were performed with ion numbers corresponding to low and high salt concentrations (4M NaCl solution). 3D models were visualized in PyMOL (The PyMOL Molecular Graphics System, Version 2.3.2 Schrödinger, LLC).

## Results and discussion

### High salt concentration leverages the enzymatic activity of PETase

Chromogenic assays using p-NPA as a substrate were conducted to explore the effect of environmental conditions on PETase enzymatic activity. To optimize the reaction conditions, reactions were supplemented with the following chemical additives: NaCl (0.1-5M), glycerol (0–25.0%), SDS (0-4.0%), and DMSO (0-100%).

Strikingly, a dramatic increase of the PETase enzymatic activity at NaCl concentrations from 500 mM to 5 M was observed (Fig. 1 *A-F*). The highest specific activity (3.8-fold increase) was measured at 5M NaCl. High activities were observed in seawater (salinity of ~35 parts per thousand (ppt), 0.47 M NaCl). In addition, increased activities in the presence of SDS <0.5% and 5% (v/v) glycerol were detected. SDS (above 3%) and DMSO negatively affected the activity.

To our knowledge, this is the first demonstration of significantly leveraged PETase activity in the presence of NaCl concentrations above 500 mM and up to 5 M. Published studies of the PETase activity were conducted at NaCl concentrations typically ranging from 0 to 150 mM (3, 11, 12, 16, 17, 25). To explain the effect of high salt concentration, a predictive analysis of conformational dynamics of PETase bound to a substrate was conducted.

### PETase/polymer complex exhibits distinct conformations at low and high salt in MDS modeling

MDS was applied to understand the parameters of PETase/substrate interactions at low and high salt concentrations. A theoretical 3-unit PET polymer model represented a hydrolytic substrate. Published 3D crystal structures of the recombinant *I. sakaiensis* PETase (a 2.02A resolution structure (PDB:6ANE) (11) and a 1.58A resolution structure of the R103G/S131A mutant PETase complex with (2-hydroxyethyl) 4-methyl terephthalate (HEMT) (PDB:5XH3) (17)) were used to build dynamic models. Covalently bound HEMT was substituted with a 3-unit PET polymer (Fig. 2*A-C*). The central unit of a 3-unit substrate was reoriented to match the positions of HEMT within the PETase catalytic center. MDS predicted the enzyme/substrate conformation states at 150 mM (low) to 4M (high) NaCl concentrations, and 0 ns to 500 ns timepoints.

**Figure 2.**
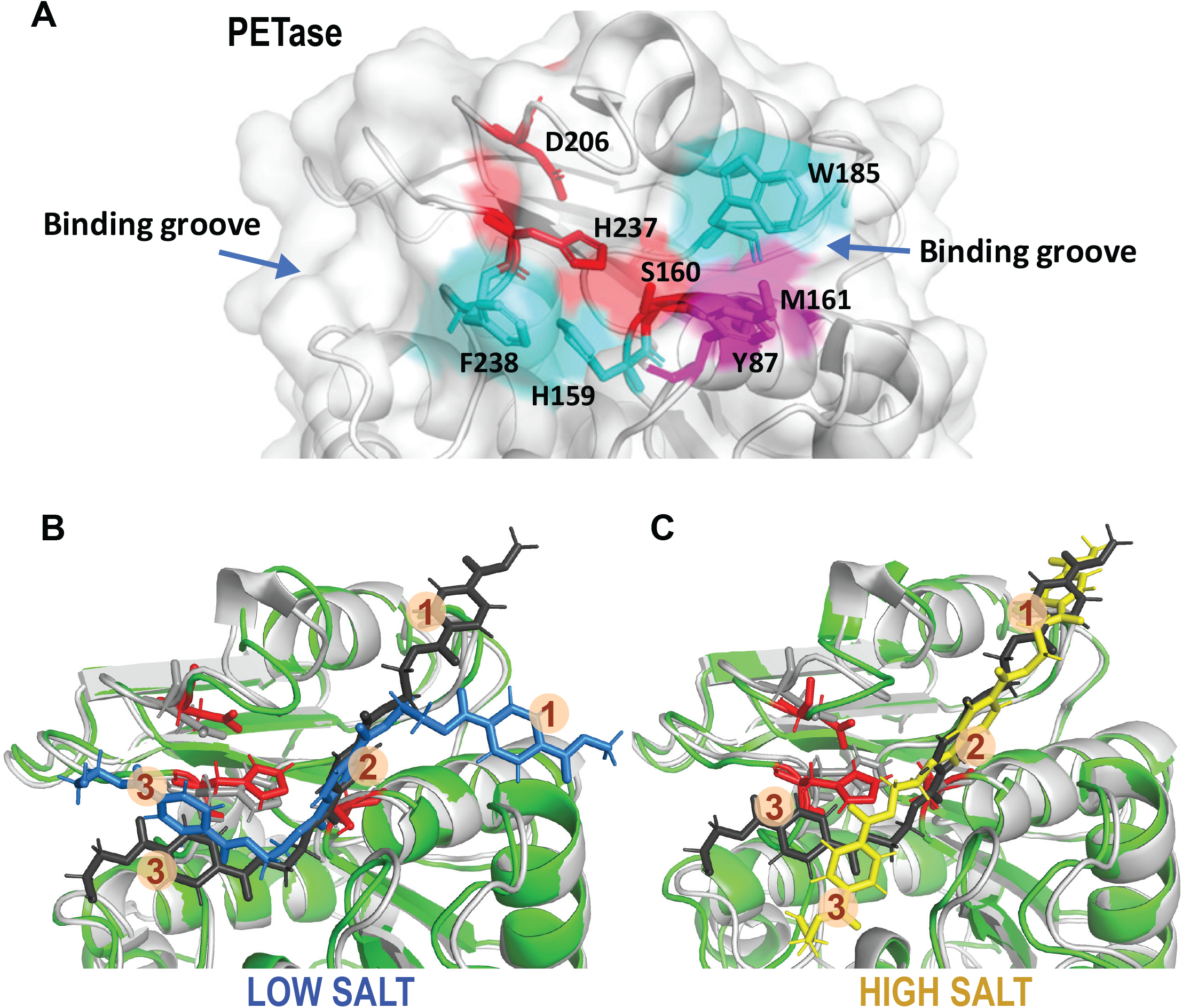
Molecular dynamics simulations predicted salt-dependent conformation changes in the PETase/polymer complex. **(A)** The 3D model is based on PDB:6ANE. Residues D206, H237, and S160 of the catalytic triad are highlighted in red. Oxyanion hole residues Y87 and M161 are highlighted in magenta. Cyan color highlights residues that influence the enzymatic activity in the PETase mutant. **(B-C)**, MDS of PETase bound to a 3-unit PET polymer. The predicted superimposed complex of PETase with a bound 3-unit PET polymer at low (B) and high (C) NaCl concentrations. Polymer units are numbered. Protein conformation states at 0 ns and 500 ns timepoints of MDS are colored gray and green, respectively. Polymer conformation states at 0 ns timepoint are shown in black. Polymer conformation states at 500 ns timepoint are shown in blue (B, low salt) or yellow (C, high salt) colors.

According to MDS, salt mediated dramatic conformational changes in the polymer-bound PETase catalytic center by altering the orientation of the polymer’s ester bonds relative to the catalytic residues (Fig. 2*B*-*C*). At low salt concentrations, the positions of units 1 and 2 and PETase catalytic residues at 500 ns simulation timepoint matched the initial geometry, and unit 1 and 3 attained bent conformations. In contrast, high NaCl concentrations mediated conformational changes in **D^206^**SIAPVNSS^214^ and I^232^NGGS**H^237^**FCANSGN^244^ catalytic loops and the polymer units relative to the catalytic residues. As a result, unit 3 of a substrate polymer became significantly extended. Notably, unit 1 retained its position near the W185 loop. The close vicinity of the catalytic triad residues D206, H237, and S160 relative to the scissile ester bond between units 2 and 3 potentially facilitated hydrolysis.

We concluded that PETase exhibits salt-dependent flexibility of its catalytic and binding centers. Modulated by environmental factors, such as salt concentration, the enzyme can fine-tune the conformation of its key residues to leverage hydrolysis of a broad spectrum of natural and artificial polymers. In turn, organisms expressing PETase-like enzymes can utilize variable carbon sources to invade new environmental habitats. Notably, the adaptability of PETase may be indispensable to advance plastics bio-utilization technologies. Ocean salt water, salt-affected soils, and locations with the increased salinity can potentially serve as efficient sites for *in-situ* bio-decomposition.

## Abbreviations

PET: polyethylene terephthalate
PETase: PET hydrolase
MD: molecular dynamics
MHET: mono-2-hydroxyethyl acid
BHET: bis-2-hydroxyethyl terephthalic acid
p-NPA: P-nitrophenyl acetate
HEMT: (2-hydroxyethyl) 4-methyl terephthalate

## Author contributions

S.A.S.: methodology, enzymatic analysis, review, editing, visualization. P.C.: software, MDS analysis, review, editing, visualization. A.V.C.: conceptualization, methodology, project administration, analysis, data curation, writing original draft, review, editing, visualization.

## Acknowledgments

The authors received no specific funding for this work. The authors thank Dr. Alex Y. Strongin for helpful discussions.

